# Supervised learning of protein thermal stability using sequence mining and distribution statistics of network centrality

**DOI:** 10.1101/777177

**Authors:** Ankit Sharma, Ganesh Bagler, Debajyoti Bera

**Affiliations:** Center for Computational Biology, Indraprastha Institute of Information Technology Delhi, Delhi, 110020, India

## Abstract

**Motivation:** It is expected that the difference in the thermal stability of mesophilic and thermophilic proteins arises, in part at least, from the differences in their molecular structures and amino acid compositions. Existing machine learning approaches for supervised classification of proteins rely on the features derived from the structural networks and the amino acid sequences. However, the network features used leave out several important network centrality values, the statistic used is a simple average and the sequence features used are hand-picked leading to an accuracy of 90%.

**Results:** We show that discriminating sub-sequences of the amino acid sequences can significantly improve classification accuracy compared to the existing approaches of counting amino acids, di-peptide or even tri-peptide bonds. We identify notions of network centrality, specifically that depends on the distances between *Cα* atoms, that appears to correlate better with thermal stability compared to the existing network features. We also show how to generate better statistics from the node- and edge-wise centrality values that more accurately captures the variations in their values for different types of proteins. These improved feature selection techniques make it possible to classify between thermophilic and mesophilic proteins with 96% accuracy and 99% area under ROC.

**Availability:** The dataset and source code used are available at https://github.com/ankits0207/Protein_Classification_BIO699

online.

## 1 Introduction

Thermal stability of a protein is a property by the virtue of which it retains its configuration when subjected to moderately high temperatures. On the basis of thermal stability, proteins are classified into two broad categories: mesophilic (<41°C) and thermophilic (80°C) proteins. Identification of factors responsible for thermal tolerance of proteins has been an active area of research and has direct implication in bioengineering of enzymes that can withstand stressful conditions like higher temperatures. There has also been significant progress in prediction of thermal tolerance using supervised and unsupervised machine learning techniques. These techniques rely on the different structures of a protein, namely, its amino acid sequence (primary structure), structure and composition of *α*-helixes and *β*-sheets (secondary structure), three-dimensional structure (tertiary structure), molecular protein volume (quaternary structure) (Szilagyi and Zavodszky (2000); *Sadeghi et al.* (2006); *Dalhus et al.* (2002); Gao and Ding (2017)). The number of possible features could easily reach hundreds or thousands which negatively impacts the time-complexity of classification algorithms. Therefore it is crucial to identify a small but effective set of features. We focus on identifying novel features from the primary and the tertiary structure in this paper.

The primary structure is usually thought to contribute most to a protein’s characteristics. Gromiha et al.(Gromiha and Suresh (2008)) used occurrences of Lys, Arg, Glu, Val and Lie to obtain 89–91% accuracy in two datasets. They reported that different machine learning methods like Bayes rule (NB), logistic regression (LR), neural networks (ANN)and support vector machines (SVM) produce comparable accuracy. Instead of counting frequencies of single amino acids, using as features the frequencies of *n*-grams (of suitable sizes) on a reduced set of amino acid alphabets was shown to be more effective with 91.8% accuracy (Albayrak and Sezerman (2012)). Recently, He et al.(He *et al.* (2016)) reported 94.44% accuracy by using frequencies of fragments (of a large size) of an amino acid sequence. Both the last results were reported for SVM. Gao et al.(Gao and Ding (2017)) conducted an extensive study in which he experimented with both amino acid frequencies as well as di-peptide frequencies on SVM, Bayesnet, ANN, and LR but their best accuracy was 86% (area under ROC was 0.91 and MCC was 0.72).

The 3D tertiary structure of a protein is generally converted to a network for the classification problem. Two kinds of network has been constructed in the past, residue interaction network (RIN) and protein structure graph (PSG). The notable difference between PSGs and RINs exist at node level as PSGs consist of amino acid residue as nodes instead of *C − α* atoms in RINs. Brinda and Vishveshwara (2005) *et. al* constructed the PSGs as a network of non-covalent interactions among amino acids thresholded by a cutoff distance. They were able to establish the importance of certain network features like hub frequency, edge count, cluster size etc. in imparting thermal stability to proteins. However, RINs are constructed with atomic level details as compared to the coarse grained representation in PSGs and are deemed to retain fine-grained structural information. RINs were employed by (Gao and Ding (2017)) who identified characteristic path length and closeness centrality as two network features which are important in imparting thermal stability to proteins. They reported 90.4% accuracy (area under ROC was 0.96 and MCC was 0.81) by using network features along with amino acid and di-peptide composition frequencies.

*Chakravorty et al.* (2017) applied numerous machine learning techniques using three different aspects of proteins: nucleotide sequence, amino-acid composition and structural configuration. They marked the importance of amino acid composition over nucleotide sequences and structural configuration as a feature to be used for classification. They reported classification accuracy of 90.97% with 10 folds cross validation. Miotto *et al.* (2018) proposed a graph-theoretic framework in which they modelled proteins as energy weighted graphs which is also built upon the concept of RINs. They used their representation to classify proteins with an accuracy of 76% and area under the ROC curve as 78%. *Ai et al.* (2018) reported that primary structures had a more profound impact on thermal stability than secondary structures. They classified mesophilic and thermophilic proteins with a classification accuracy of 84.07%.

The motivation behind this paper was to explore new sequence and structure based features which could be used to distinguish between mesophilic and thermophilic proteins. For incorporating feature values we argue that the use of average statistics of network parameters misrepresents the heterogeneity across residues. We propose two types of distribution statistics to capture the fine-grained variations. Existing studies reveal that thermal stability is manifested as robustness of nodes in RINs, and thus, we explore network features that arise from network centrality and inter-node distances. Further, unlike previous studies which have used fixed-length sequence-based patterns, we identified discriminating sequence features of all possible length. By investigating a set of 575 thermophilic and 2032 mesophilic proteins, we built classifiers to achieve an overall F1 score as 0.96, area under ROC curve as 0.99 and MCC as 0.91.

## 2 Materials and Methods

As is the case with any supervised learning experiment, we started by collecting data, followed it by extracting features and finally applying various supervised learning techniques.

### 2.1 Data preparation

Despite extensive research articles published on protein thermal stability classification and prediction, most of the datasets that were used in the corresponding studies were not available at the time we undertook this study. Therefore we started by preparing and labelled-dataset of thermophilic and mesophilic proteins that we have also made freely available for anyone to download.

We started with a list of 94 mesophilic and 9 thermophilic organisms (See Supplementary Data 1 Takemoto *et al.* (2007)) and compiled a list of 3386 mesophilic proteins and 716 thermophilic proteins belonging to these organisms using the Protein Data Bank (Berman *et al.* (2006)) — the proteins themselves were selected and classified using their optimal growth temperatures (Takemoto *et al.* (2007)). The resolution of the protein structures thus obtained varied between 0.93 Å and 19.8 Å. We further pruned this list and retained proteins with resolution at most 2.5 Å since a high resolution implies greater error in capturing the exact coordinates of amino acid residues.

This gave us 598 thermophilic and 2578 mesophilic proteins; however, when we plotted the number of proteins against resolution (see Fig 1) we observed that this dataset has a larger number of mesophilic proteins with high resolution as compared to thermophilic proteins. Higher resolution indicates a larger error in capturing the exact coordinates of the amino acid residues. To avoid the bias against mesophilic proteins we undertook further refinement steps. In the presence of multiple structures for the same protein, the structure with a higher molecular weight was considered assuming it to be richer in terms of captured residues. In case of multiple proteins after this step, the structure with the lowest resolution was kept. After all these steps, we obtained 575 thermophilic and 2032 mesophilic proteins with the best resolution (See Supplementary Data 2 & Supplementary Data 3). Fig 1 shows the distribution of structures in the curated data set against their resolutions; it can be observed that the distribution is almost identical between thermophilic and mesophilic proteins which makes the dataset ready for feature extraction and learning.

**Fig. 1.**
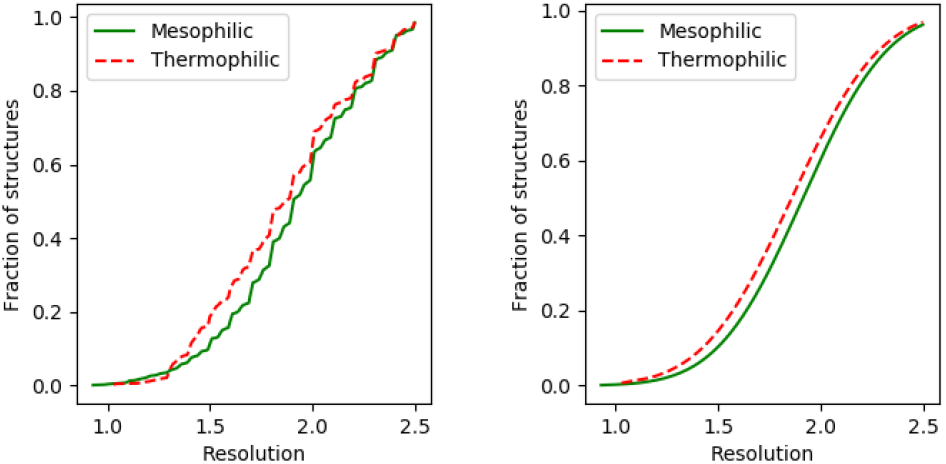
Normalized cumulative distribution of the number of proteins against their resolutions (Å). Left chart shows the unrefined dataset with 598 thermophilic and 2578 mesophilic proteins and right chart shows the final dataset with 575 thermophilic and 2032 mesophilic proteins.

### 2.2 Structure-based features (NF)

Structural compactness has been known to be one of the critical features of thermal stability (Robinson-Rechavi *et al.* (2006)). For identifying the important structural features, every protein was modeled as a weighted UN-directed “residue interaction network” (RIN) that is a network of the constituent amino acids which are interconnected depending on their spatial proximity. The *Cα* atom of an amino acid was used as its representative node in the RIN and any two such nodes were linked (with a weighted edge) if they were within a distance of 6.5 Å(Gao and Ding (2017)). The difference between the actual distance between the amino acids and this threshold was used as the weight of the link between them. Thus the weight decreased as the amino acids went further apart, thereby weakening their interaction.

Previous studies have identified network properties of the RINs such as degree, number of edges, largest cluster size, weighted clustering coefficient, etc. that are deemed important for thermal properties of proteins (Brinda and Vishveshwara (2005); Gao and Ding (2017)). We noticed that many of these are properties that fail to assess the contribution of a node to the rigidity of a network. This motivated us to explore network centrality of nodes as network features. A node with high centrality is understood to play a central role in holding a network together. As usual there can be different notions of relative importance in a work giving rise to several centrality measures. Below we review the network centrality measures that we used in our experiments. In addition to the network centrality measures, we also used characteristic path length and weighted clustering coefficient; even though these are not centrality measures, these two network features were recently reported (Gao and Ding (2017)) so we wanted to evaluate how they match up against the other centrality measures.

A list of all the network features evaluated by us are presented in Table 1. The notations used below are summarized in Table 2.

**Table 1.**
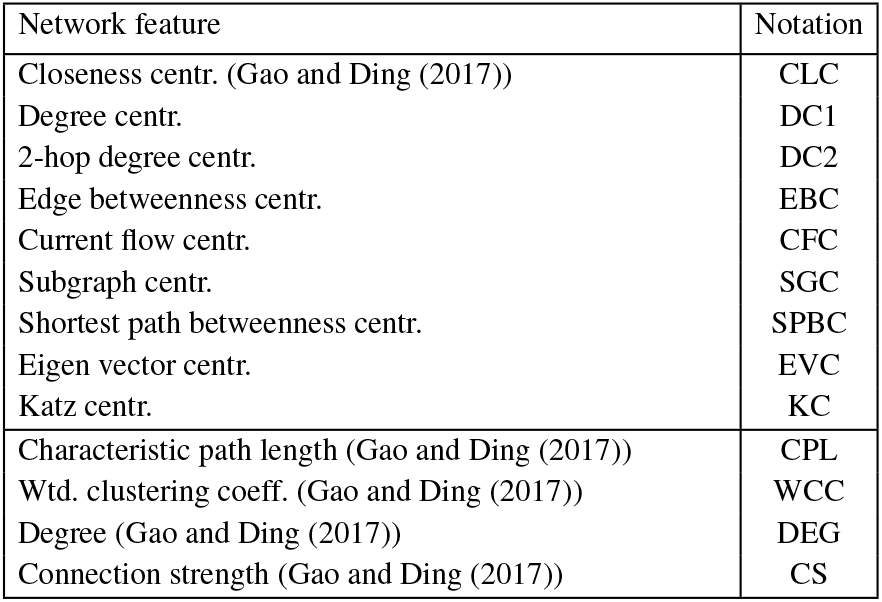
Network features used for supervised learning of thermal stability. All but the last four features are centrality notions.

| Network feature | Notation |
| --- | --- |
| Closeness centr. (Gao and Ding (2017)) | CLC |
| Degree centr. | DC1 |
| 2-hop degree centr. | DC2 |
| Edge betweenness centr. | EBC |
| Current flow centr. | CFC |
| Subgraph centr. | SGC |
| Shortest path betweenness centr. | SPBC |
| Eigen vector centr. | EVC |
| Katz centr. | KC |
| Characteristic path length (Gao and Ding (2017)) | CPL |
| Wtd. clustering coeff. (Gao and Ding (2017)) | WCC |
| Degree (Gao and Ding (2017)) | DEG |
| Connection strength (Gao and Ding (2017)) | CS |

**Table 2.**
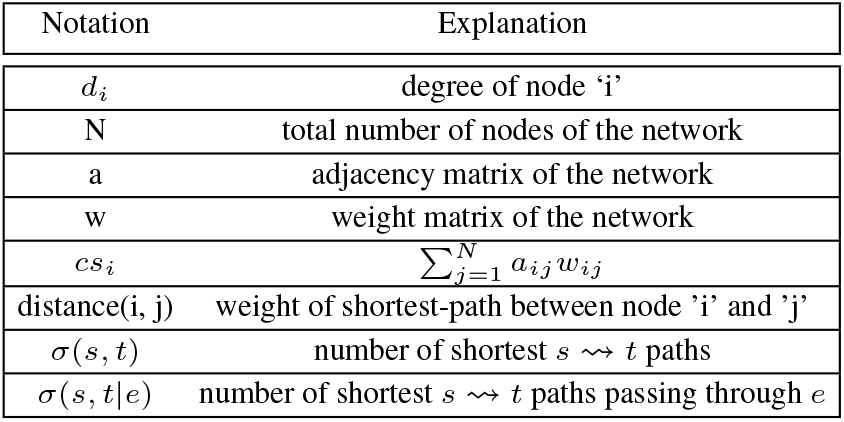
Notations associated with structure-based features

1. **Closeness Centrality (CLC)**: The closeness centrality of a node corresponds to the nearness of all nodes in a network to that node. For a node *i* it is defined as 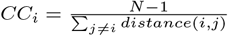.
2. **1-Hop degree centrality (DC1)**: The degree centrality of a node in a network is defined as the fraction of nodes reachable from that node by one edge.
3. **2-Hop Degree Centrality (DC2)**: The 2-hop degree centrality is an extension of the 1-hop degree centrality. It is defined as the fraction of nodes that are within 2 hops of a node.
4. **Edge Betweenness Centrality (EBC)**: The edge-betweenness centrality of an edge is defined as the sum of the fraction of all-pairs shortest paths that pass through that edge (Brandes (2001)). Mathematically, edge betweenness centrality of an edge *e* is expressed as 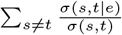.
5. **Current-flow Closeness Centrality (CFC):** Current-flow closeness centrality, also known as *information centrality*, (Brandes and Fleischer (2005)) is variant of closeness centrality based on effective resistance between nodes in a network. Mathematically, current flow closeness centrality of a node 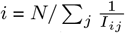, where the matrix *I*_*ij*_ is computed from two matrices *D* (a diagonal matrix with the degree of each node along the diagonal) and *J* (a matrix with all elements equal to 1) as 1/*I*_*ij*_ = (*B^−^*^1^)_*ii*_ + (*B*^−1^)_*jj*_ − 2(*B*^−1^)_*ij*_ in which *B* = *D* − *A* + *J*.
6. **Subgraph Centrality (SGC)**: Subgraph centrality of a node *i* is defined as the sum of the weights of all weighted “closed walks” of all lengths starting and ending at node *i* (Estrada and Rodriguez-Velazquez (2005)). Computed using a spectral decomposition of *A*, it is expressed as 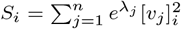 where *v*_*j*_ is an eigen vector of the adjacency matrix *A* corresponding to the eigen value *λ*_*j*_.
7. **Shortest Path Betweenness Centrality (SPBC)**: The (node) betweenness centrality of a node is defined as the fraction of shortest paths, aggregated over all pairs of source and destination nodes, that pass through a node. It is similar in nature to EBC.
8. **Eigen-vector Centrality (EVC)**: Eigen-vector centrality is defined using an eigen-vector of the adjacency matrix representing a network. If *x* denotes one such eigen-vector, then the eigen-vector centrality of the *k*-th node is the *k*-th attribute of *x*.
9. **Katz Centrality (KC)**: Katz centrality is a generalization of eigenvalue centrality, and for the *k*-th node it is defined as *x*_*i*_ = *α* Σ_*j*_ *A*_*ij*_ *x*_*j*_ + *b*, where *A* denotes the adjacency matrix.

In addition to **DC1** and **DC2** we also tried 3-hop degree centrality (the fraction of nodes reachable within 3 hops) but found it to be a poor classifier and hence it was not considered. In our experiments we decided to evaluate the above features along with a few non-centrality network features the first two of which are closely related to **DC1**.

1. **Degree (DEG)**: The degree of a node (representing the *Cα* atoms) is defined as the number of edges incident upon it (representing the number of other *Cα* within the threshold distance). This is essentially **DC1** multiplied by the number of nodes in a network.
2. **Connection Strength (CS)**: The connection strength of a node is defined as the total weight of the edges incident on that node. This can be seen as a weighted variant of **DEG**.
3. **Characteristic Path Length (CPL)**: The characteristic path length of a network is defined as the average length of the shortest paths between all the pair of nodes,i.e., Σ_*i≠j*_ *distance*(*i, j*)/*N* (*N* − 1). A small CPL indicates that most nodes are close to each other and points to a tightly-knit structure. We decided to evaluate this alongside centrality measures since it is closely related to **CLC**.
4. **Weighted Clustering Coefficient (WCC)**: Clustering has been repeatedly found to be responsible for increased thermostability (Brinda and Vishveshwara (2005); Vijayabaskar and Vishveshwara (2010)); so we decided to include weighted clustering coefficients in our experiments that are well-established measures of clustering. The WCC of a network measures the degree to which the nodes of the network tend to cluster together. It counts the number of triangles that a node participates in, along with normalization and taking into consideration the weights of the triangles. We used the following formula to calculate it.

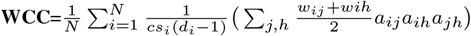

For identifying important features among the thirteen listed above, we first performed feature selection using 10-fold cross validation. We observed that the features **SPBC**, **EVC** and **KC** were not consistent across all folds and hence they were dropped. Further we found that seven features, those listed in Table 3, were present in almost all folds. Next we experimented by holding-out each of the remaining features individually and once again those seven features yielded the largest drops in F1 score, area under ROC curve and MCC. We decided to constitute our final set of network-structure based features from these seven features (more so since we were interested in the role of centrality metrics) and we refer to this set as **NF**. The results of the hold-out experiments for different supervised learning frameworks are presented in Supplementary Data 1 in Tables S1 (random forest), S2 (naive Bayes), S3 (support vector machine) and S4 (artificial neural network).

**Table 3.**
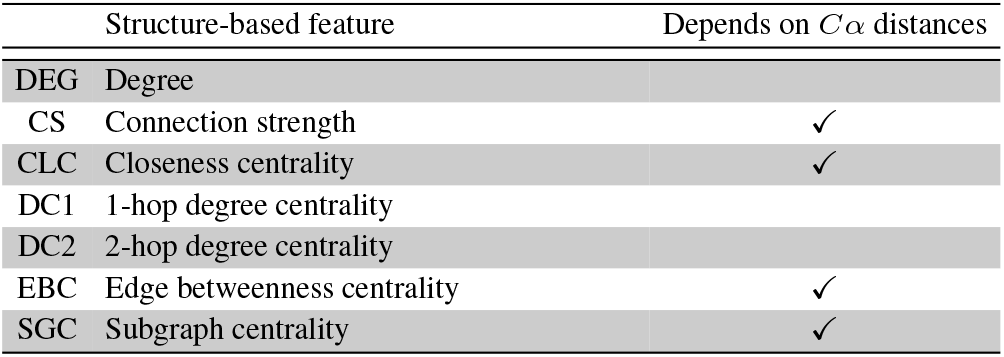
Final list of network-structure based features (denoted **NF**). The ones that are ticked are computed using the distances between *Cα* atoms.

### 2.3 Distribution statistics of network feature values

We discussed in detail an exhaustive set of network features in the previous subsection that we used for identifying structural differences between mesophilic and thermophilic proteins. Some of the these features involve computing a value for every node (e.g., betweenness centrality) or every edge (e.g., edge betweenness centrality). In the existing approaches of machine learning classifiers, these values are incorporated in the form of their average values (Gao and Ding (2017)).

We suspect that average values are not suitable for capturing the subtle differences between the structures and conjecture that the distribution of node-specific (or, edge-specific, for the edge-based features) values can provide valuable insight. For example, the average EBC values of both the mesophilic protein 2WH7 and the thermophilic protein 1MTP are close to 0.006. However, almost all EBC values in the latter lie in the narrow range of 0 to 0.07 whereas the same in the former protein spreads across a wider range of 0 to 0.5 (see Fig.2(B)).

**Fig. 2.**
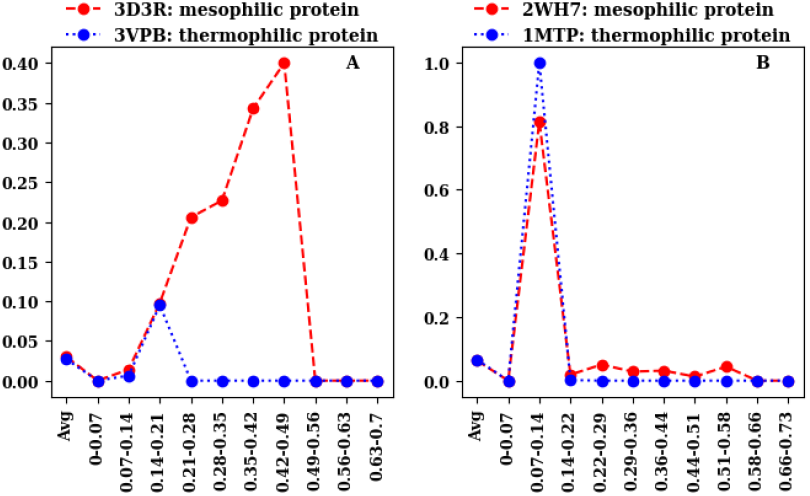
Examples illustrating that use of average value of features (EBC in this case) is not a good approach as proteins belonging to different thermal classes may have similar average values but the distribution across bins can vary which could be exploited for classification. The *X*-axis represents the various bins used for the EBC values and the *Y*-axis represents the distribution statistic, DSA in Fig. (A) and DSF in Fig. (B). The distribution statistics for both pairs of proteins are evidently different despite the fact that the average EBC values (plotted as **Avg**) are identical.

Hence we proposed two different statistical summary to incorporate the distribution of centrality values in a fine-grained manner in our classification features. The existing approach of using the average of all the values is referred to as **AVG**.

Consider a network feature that initially computes values at every node (or at every edge). The methods below generate an array of summary values from these individual values.

1. First we compute the value at all node (or at all edge).
2. The minimum and maximum values are extracted amongst all the nodes.
3. The range in between the maximum and minimum values was split into a fixed number of equally spaced non-overlapping bins. We chose 10 bins after visually inspecting the ranges of values for our network features.
4. Each bin is populated with the nodes (or edges) having value that lies in the range of that bin.
5. The summary array is generated from these bins.
  - In **frequency distribution statistic (DSF)** an array containing the *fraction of nodes in every bin* (normalized by the total number of nodes in the entire network) is constructed.
  - In **average-value distribution statistic (DSA)** an array containing the *average value in every bin* (normalized by the total number of nodes in the entire network) is constructed.
6. These arrays are returned to be used as features for supervised classification.

In the AVG statistic a feature vector would have only a single value representing the cumulative average, whereas in DSF and DSA a feature would have 10 values, one for each bin; for example, for DSF these values correspond to the fraction of *Cα* atoms whose network feature values (or sequence feature values) fall in a particular bin. Our hypothesis is that the centrality values for the thermophilic proteins exhibit a more compact behaviour compared to that of the mesophilic proteins, and therefore, the DSF values are concentrated in fewer bins for the former as compared to the latter.

### 2.4 Sequence-based features (SF)

The primary structure of a protein is a linear sequence of amino acids. This sequence is naturally the obvious target to probe for structural and functional differences. Indeed, the use of sequence information to analyse thermal behaviour is a quite common approach and follows two main themes (Haney *et al.* (1999b); Chakravorty *et al.* (2017); Gao and Ding (2017)).

1. **Amino acid composition (AAC):** Amino acid composition of a protein refers to frequencies of each of 20 amino acids (normalized with respect to the total number of amino acids in the protein) that appear in a protein. This is the most common sequence feature of proteins that is used for thermal stability analysis (*Chakravorty et al.* (2017), Gao and Ding (2017)). We optimized further by only considering the normalized frequencies of Alanine (A), Glutamine (Q), Glutamic acid (E), Histidine (H), Threonine (T). These five amino acids were chosen since they are known to have a significant role in thermal stability (Farias and Christina Manhães Bonato (2002); Ebrahimi *et al.* (2011); Haney *et al.* (1999a)) and they presented a huge improvement in classification performance even in our experiments (see Fig. S1 in Supplementary Information).
2. **Di-peptide composition (DP):** Similar to **AAC**, in a di-peptide composition of a protein we store the number of di-peptides of each type in a protein, normalized by the total number of di-peptides in the protein (Gao and Ding (2017)). The role of di-peptides in thermal stability is also well-established; in fact, Ebrahimi *et al.* (2011) was able to predict thermostability by only considering amino acid and di-peptide frequencies. We added to this list two more sequence features that we now explain.
3. **Tri-peptide composition (TP):** Noting the importance of peptide bonds, we stored the number of each tri-peptide in a protein, normalized by the total number of tri-peptides in the protein. To the best of our knowledge, the role of tri-peptides for thermal stability has not been investigated. We did not continue further to store 4 and more peptide bonds since our experiments did not reveal any significant improvement in classification quality.
4. **Discriminating Sequential Patterns (SP):** Unlike the previous three sequence features that count the frequency bonds of specific lengths (one, two or three), we propose to first identify the peptide chains (of any length) that frequently appear in one class of proteins and infrequently in another class. We call these discriminating sequences. For finding the discriminating sequences we leverage results from the sequential mining literature that identify substrings in a sequence of symbols with support larger than a threshold. Support of a substring *ϕ* in a database of sequences is the total number of times *ϕ* appears in the entire database; note that a substring may appear multiple times in a sequence and this multiplicity is taken into account. Also, substrings must appear *contiguously*, i.e., “ABC” is a substring of “XYABCABC” with support 2 but not a substring of “XAYBC”. Fischer et al. (Fischer *et al.* (2008)) introduced the problem of *emerging substring mining* whose objective is to find all substrings *ϕ* from two databases of sequences 𝒟_1_ and 𝒟_2_ such that *support*(*ϕ*, 𝒟_1_) *≥ P*_*s*_ and *support*(*ϕ*, 𝒟_2_) *≤ support*(*ϕ*, 𝒟_1_)/*P*_*g*_; the parameter *P*_*s*_ indicates that *ϕ* must appear abundantly in 𝒟_1_ and *P*_*g*_ (satisfying *P*_*g*_ > 1) indicates that *ϕ* appears in 𝒟_2_ infrequently. The emerging substrings are a natural candidates for differentiating between the databases 𝒟_1_ and 𝒟_2_. So we first identify the emerging substrings of amino acids that are relatively more frequent in mesophilic compared to thermophilic or relatively more frequent in thermophilic compared to mesophilic using the algorithm of Fischer et al. Then we use the support values of these substrings as features for classification. We performed an exhaustive search over the space of tunable parameters *P*_*s*_ and *P*_*g*_. *P*_*s*_ = 0.45 and *P*_*g*_ = 1.7 were chosen as the optimal parameters based on the number of substrings it generates and the classification performance. These substrings constituted 17 di-peptides and 7 tri-peptides. By varying the parameters we could obtain quad-peptides too but the list of substrings in those cases was significantly large. We wanted a small list of emerging substrings that yield good classification performance and hence we finalized those 24 as SP; the entire list of SP along with AAC are presented in Supplementary information (Table S8).

### 2.5 Supervised learning classifiers

For supervised classification we used four classifiers of different nature that we explain below.

#### Naive Bayes classifier (NB)

Naive Bayes classifier is a probabilistic machine learning classifier that works based on the principle of Bayes theorem. It is a simple classifier, converges faster and works well even with lesser data.

#### Support Vector Machine (SVM)

Support vector machines belong to the class of supervised learning algorithms in which the aim is to identify a hyper-plane which separates the data belonging to different classes. SVM was chosen as one of the potential classifiers to be used for classification because it can be efficiently scaled with high dimensional data which is obtained due to binned feature extraction. The hyper parameter setting involved the use of ‘rbf’ kernel with probability estimates set to true.

#### Artificial Neural Network (ANN)

Artificial neural network is a supervised learning framework which progressively improves the prediction by the virtue of back-propagation in which the weights are readjusted after computing the difference in between the ground truth and the prediction. ANN with ‘relu’ as the activation function and ‘adam’ as the stochastic gradient descent solver was used.

#### Random Forest (RF)

Random forest is an ensemble supervised learning technique which runs over a collection of decision trees and makes prediction on the basis of consensus. Random forest works on the principle of traversing over random subsets of features in search of the best feature. This imparts diversity to the search space and generally results in a robust prediction. Being a tree based ensemble method, this is a popularly chosen model as it can handle the problem of imbalance amongst classes.

### 2.6 Evaluation metrics

The dataset that we curated (explained in Subsection 2.1) contained only 575 thermophilic but 2032 mesophilic proteins. To ensure that learning isn’t biased due to imbalance in data, down-sampling was used to create 4 smaller datasets of 500 mesophilic proteins; all the 575 thermophilic samples were retained. We report the average of 10-fold cross validation on these datasets, further averaged over the 4 datasets. For performance evaluation we used the standard metrics of F1-score, area under the ROC curve and Matthews correlation coefficient (MCC).

## 3 Results

Towards the objective of creating machine learning models for classification of thermophilic and mesophilic proteins, we investigated network-based features extracted from residue interaction graph models and those obtained from amino acid sequences. A total of seven network-based features (CLC, EBC, DC1, DC2, CS, SGC, DEG) capturing various structural aspects critical for their thermal response were computed. Similarly, four sequence-based features enumerating amino acid count (AAC), di- and tri-peptide composition (DP and TP), and frequently occurring sub-sequence patterns (SP) were used for building models. Importantly, we capture the nature of distribution of node- and edge-centric network properties that are often ignored if their average values are used.

### 3.1 Distribution Statistics

The success of classification models is critically dependent on the use of appropriate features that reflect the distinguishing aspects. With respect to the network features, capturing them at the correct level of granularity is vital. We surmise that using average values of network features yield moderate classification accuracy across different classifiers. The nature of variation in the network properties is lost in the process of averaging (Figure 2). Hence, we computed distribution statistics by binning each of the network feature to measure the frequency (DSF) and average (DSA) values. We observe that performance of classification improves by using the distribution statistics (Figure 3). Compared to the model implementing averaged values, DSF and DSA fared better using SVM, ANN and RF. Random Forests emerged as the best classifiers in terms of F1-score, AUC and MCC scores, yielding an F1 score of 0.87, AUC of 0.93 and MCC of 0.74 when DSA was used and an F1 score of 0.88, AUC of 0.94 and MCC of 0.75 when DSF was used. Being a weak classifier, the performance of Naive Bayes when measured using F1 score was best when cumulative average was used instead of any distribution statistic. However, even NB lagged behind the other two when AUC and MCC scores were compared. This trend clearly validates our hypothesis that an extremely coarse statistic like cumulative averaging does not capture the fine-grained variations in the values. The bins used by us are ad-hoc in nature but could be learned from the data using, e.g., clustering and supervised learning; however, we leave that for future since DSA and DSF outperformed AVG quite handsomely.

**Fig. 3.**
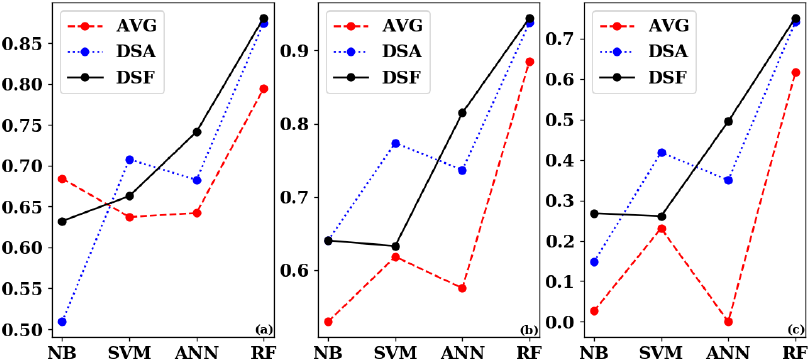
Comparison of distribution statistics for proposed NFs using different classifiers: (a) F1 score (b) Area under ROC curve (c) MCC.

For the next set of experiments we decided to use the frequency of each bin as features (DSF) since it gave marginally better scores for ANN and RF on all three metrics.

### 3.2 Network-based Features

We evaluated seven network features (listed in Table 3) that enumerate various structural aspects for their contribution towards classification performance. These features appeared in almost all folds of a feature selection step. However, it was to be seen whether they really complement each other for the task of classification. For this we plotted a heat-map in which we considered all pairs of features. For each pair we separately calculated their correlation coefficient for thermophilic and mesophilic proteins. Then we created a heat-map showing the difference (in absolute sense) of their correlation coefficients (see Fig. 4). We observed that there was a difference of at least 2.5% between the correlations for the two classes of proteins and the difference between some pairs, e.g., degree and closeness centrality, were higher than 50%. Surprisingly enough, there was a difference between degree and DC1 even though both depend upon the number of neighbors of a node; this justified the inclusion of both of them in our list of features.

**Fig. 4.**
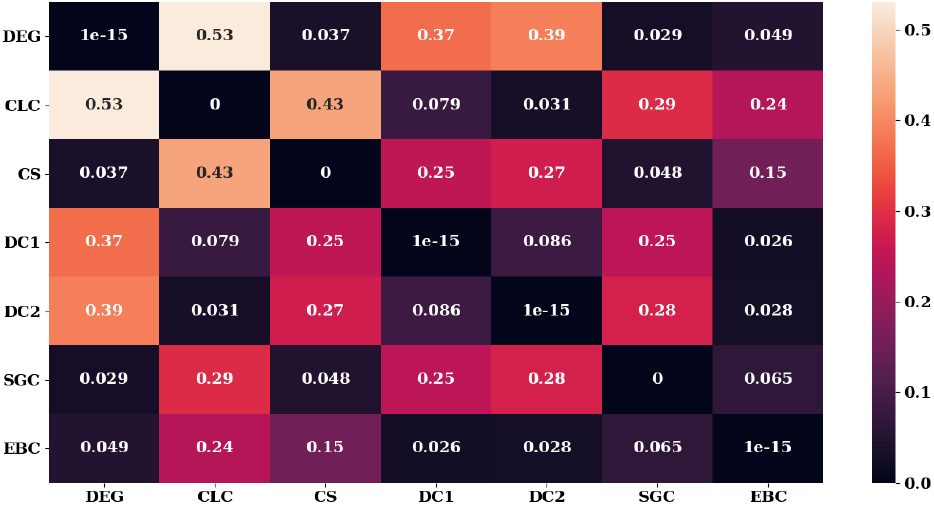
Difference in the correlation coefficient values of pairs of network features between the two classes of proteins. The darker the color, the closer are the correlation values. A very low difference for a pair of features suggests that both of them need not be considered as features. The minimum correlation is 2.5% (between EBC and DC1) and the maximum correlation is 53% (between CLC and DEG).

Table 4 shows classification performance of hold-out experiments to identify the network features that contribute the most. These held-out results are shown for the Random Forest classifier using DSF. Supplementary Information provides results of held-out experiments conducted for ANN (Table S7), SVM (Table S6) and NB (Table S5). The features in NF extract various aspects of compactness and rigidity of the protein structures through graph theoretical metrics. The results in those tables indicate that each of them contribute equally and in some unique manner. Unlike the earlier experiments reported in literature where average values were used, here we are using the distribution of the EBC, SGC, etc. values to differentiate; hence, it would not be correct to say that proteins of a particular class have high EBC or low SGC. On the contrary, our finding is purely a supervised learning observation that proteins of a particular class exhibit a significantly different pattern of centrality values compared to those of the other class.

**Table 4.** Hold out feature comparison for network features using RF. The features reported are sorted in descending order of their evaluation metric so as to highlight their contribution to the classification. The ones that are in **bold** are the novel network features employed by us for thermal stability. The ones that are <u>underlined</u> are based on edge-weights.

| Experimental setup | F1 score | Area under ROC | MCC |
| --- | --- | --- | --- |
| **All NFs** | 0.90 | 0.95 | 0.76 |
| Held-out feature |  |  |  |
| **<u>SGC</u>** | 0.75 | 0.86 | 0.69 |
| **<u>EBC</u>** | 0.75 | 0.86 | 0.69 |
| **<u>CS</u>** | 0.75 | 0.86 | 0.68 |
| DC1 | 0.74 | 0.86 | 0.68 |
| DEG | 0.74 | 0.86 | 0.67 |
| **<u>CLC</u>** | 0.74 | 0.85 | 0.67 |
| **<u>DC2</u>** | 0.73 | 0.85 | 0.66 |

It is evident from Table 4 (and tables S7, S6, S5 in Supplementary Information) that complex node- or edge-based centrality measures such as EBC, SGC and CLC add more value to prediction of thermal stability compared to network-wide average values like WCC and CPL. We also observed that network properties that depend upon edge-weights, and hence depend upon the distances between *Cα* atoms, are found to be equally important, if not more, as compared to properties that only depend upon the unweighted connections.

### 3.3 Sequence-based features and Composite Model

We demonstrated above how to involve centrality of RINs in a novel manner for classification. Now we discuss how sequence-based features can further improve this classification. For this we experimented by combining different sets of sequence features among AAC, SP, DP and TP. We observed that count of the amino acids Ala, Gln, Glu, His, Thr (AAC) consistently performed well, and thus, was chosen as a key component in all combinations.

Fig. 5 shows the comparison of various combinations using the RF model. Supplementary information illustrates the results of the same comparison for ANN (Fig. S4), SVM (Fig. S3) and NB (Fig. S2). We observed that AAC+DP performed slightly better than AAC+TP and much better when compared to AAC+DP+TP. This can attributed to the fact that combining DP and TP generates way too many redundant features invariably adding to noise. Therefore, it is critical to choose sequence features so as to capture the relevant information content hidden within the sequences without complicating the feature vector. This balance is achieved by using Sequential Patterns (SP) which identifies discriminating amino acid sequences of different lengths. These patterns have the ability to simultaneously find the important ones and remove the redundant ones. The 24 amino acid sequences that were obtained from SP, along with 5 amino acids (part of AAC), are listed in Table 5. It can be observed that this list contains tri-peptides too but comparatively many di-peptides. This supports the existing practices of considering di-peptides for classification (Gao and Ding (2017); Lin and Chen (2011); Zhang and Fang (2006)). Furthermore, we also listed in the table the contribution of each of the 29 sequences during the RF classification (a remarkable feature of decision tree based methods). Interestingly we get significant contributions from subsequences of all sizes, namely, amino acids, di-peptides and tri-peptides; this shows the efficacy of our proposal to employ SP for classification. Indeed, without the network features, AAC & SP combined gives the best classification, even when compared to AAC and SP individually. Classification performance improves even further when combined with NF.

**Table 5.** Amino acid sequences whose frequencies in the thermophilic and the mesophilic proteins differ significantly

|  |  |  |  |  |  |  |  |
| --- | --- | --- | --- | --- | --- | --- | --- |
| AAA | AEE | DAL | DEI | DII | LRE | LVL | FI |
| FR | GM | HG | HP | II | DN | EF | MD |
| MF | MN | NE | NR | NV | NY | RM | YI |

**Fig. 5.**
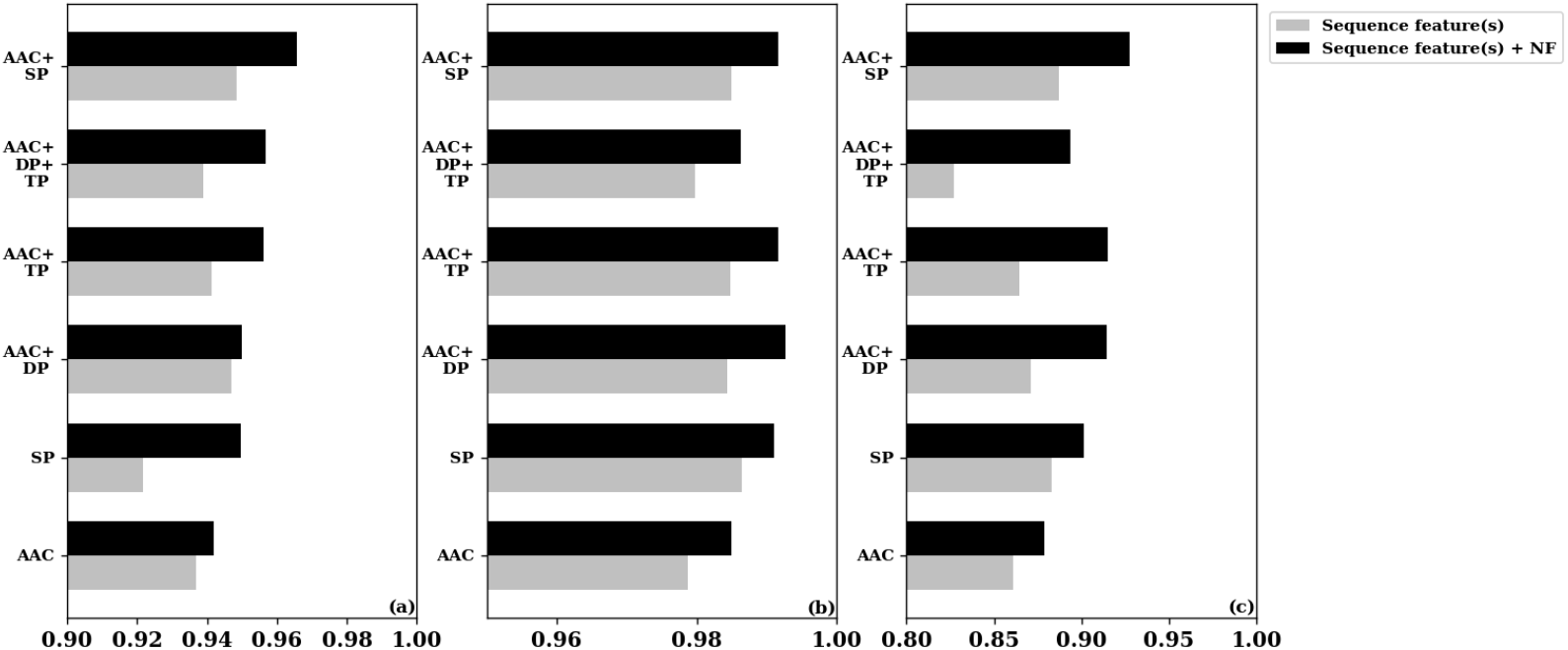
Comparison of classification using proposed network and sequence features against sequence features alone. Random forest (RF) classifier is used. The F1 score is reported in (a), area under the ROC curve is reported in (b) and MCC is reported in (c).When only network features are used, F1 score is 0.90, area under ROC is 0.95 and MCC is 0.76 (Table 4).

Based on the above experiments, we finally propose for classification the network features in NF represeted using frequency distribution statistic (DSF) and the AAC and SP sequence features. We implemented these features in RF, SVM, NB and ANN whose results are illustrated in Fig. 6. Our observation is that the combination of DSF, AAC and SP presents the best classification performance when used in the Random Forests model (F1-score=0.96, AUC-ROC=0.99, MCC=0.92, Accuracy=0.96). A further advantage of RF over ANN is that the former requires smaller training time.

**Fig. 6.**
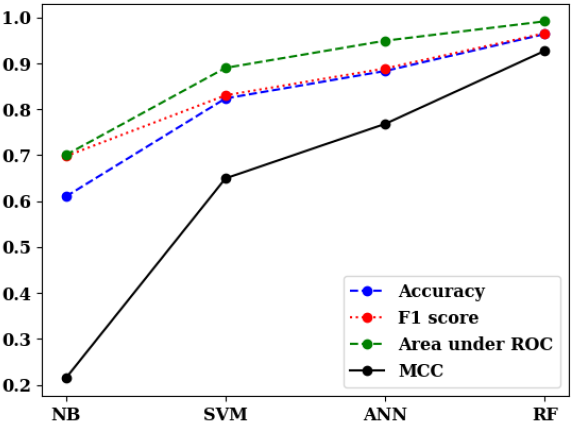
Classification performance for our proposed feature set: NF (using DSF), AAC and SP under different machine learning algorithms.

## Discussion and Conclusions

We started this work with the goal of designing a supervised classification pipeline to identify between thermophilic and mesophilic proteins based on their primary and tertiary structures. A major challenge that we faced during this work was the unavailability of a publicly available data set comprising of labelled mesophilic and thermophilic proteins. A major outcome of this work is the release of our dataset (along with the associated scripts) in public domain. The dataset that we obtained has a lesser number of thermophilic proteins as compared to mesophilic proteins (~1:4 ratio), so we had to apply preprocessing techniques to ensure that we do not introduce bias in our results for the mesophilic class.

The major outcome of this work is a conglomeration of ideas for deciding suitable features that can help in deciding thermal property of proteins. The first is the adoption of RF over other machine learning models. Recent approaches have reported 86-90% cross-validation accuracy (Gao and Ding (2017); Ebrahimi *et al.* (2011)) using SVM and using simpler network and/or sequence features. We also observed that SVM was leading over models when we used a subset of NF and our sequence features (Sharma (2018)); however, as we moved towards an enriched feature set, the performance of SVM did not improve much (accuracy was 0.82) while others, like ANN and RF, leapt forward. This is expected for SVM and similar linear models as features become more and more complex and inter-dependent.

The second idea is that of using an enriched set of edge-based network centrality properties as features. Centrality of a node captures the “importance” of a node in a network and we show that RINs of thermophilic and mesophilic proteins have nodes whose centrality distributions are quite distinct. We also focus on network properties that specifically consider the inter-atomic distances in the tertiary structure. We also explained a method to use the “distribution” of centrality (and other node/edge-specific) values as a feature in place of the average value. There could be better methods of embedding these distributions into feature space and that is left as a future research direction. The objective of this work was to only identify important centrality features but an important question that remains unanswered is the exact nature of difference between the distributions for thermophilic and mesophilic proteins and how that affect thermal stability.

The final idea is that of determining amino acid sequences whose appearances in thermophilic and mesophilic proteins are starkly different. This is quite different compared to only using di-peptides (Gao and Ding (2017); *Ebrahimi et al.* (2011)) or tri-peptides or pentamers (*He et al.* (2016)) based on heuristics. The emerging amino acid sequences that we used for classification not only has di-peptides but also tri-peptides whose role in thermal stability or other protein functions are not at all understood. Our experiments point to specific tri-peptides Ala-Ala-Ala, Asp-Glu-Ile, etc. (see Table 6) that should be studied further.

**Table 6.** 29 sequence-based features and their importance under RF.

| Top 5 amino acid sequences | Importance |
| --- | --- |
| gln (Q) | 13.37% |
| ile-ile (II) | 11.13% |
| ala (A) | 5.82% |
| met-asp (MD) | 5.78% |
| asp-glu-ile (DEI) | 5.58% |
| Rest 24 amino acid sequences | 58% |
| AAA AEE DAL DII LRE LVL FI |  |
| YI FR GM HG HP DN EF NY |  |
| RM MF MN NE NR NV E H T |  |
| Contribution of A,E,H,Q,T | 31% |
| Contribution of 17 di-peptides | 48% |
| Contribution of 7 tri-peptides | 21% |

Lastly, many stages of our classification pipeline could be applied to different functions of proteins or other biological problems, e.g., classification of cancer patients. The effectiveness and fine-tuning is a question that remains to be answered.

## Acknowledgements

This work has been partially supported by the Department of Biotechnology, India.

