## Supplement tables and figures. for "Supervised learning of protein thermal stability using sequence mining and distribution statistics of network centrality"

**Supplementary Material**

#### S1 Hold-out evaluation among all network features

| Experimental setup | F1 score | Area under ROC | MCC |
|---|---|---|---|
| **NF** | 0.90 | 0.95 | 0.76 |
| **NF** + WCC + CPL + CFC | 0.86 | 0.94 | 0.74 |
| Held-out feature | | | |
| CLC | 0.70 | 0.87 | 0.68 |
| EBC | 0.71 | 0.87 | 0.69 |
| DC2 | 0.72 | 0.87 | 0.70 |
| CS | 0.72 | 0.88 | 0.70 |
| SGC | 0.72 | 0.88 | 0.70 |
| DEG | 0.72 | 0.88 | 0.70 |
| DC1 | 0.73 | 0.88 | 0.71 |
| WCC | 0.73 | 0.88 | 0.71 |
| CPL | 0.74 | 0.88 | 0.72 |
| CFC | 0.74 | 0.89 | 0.72 |

Table S1: Network feature selection using RF among WCC, CPL, CFC and those in NF

| Experimental setup | F1 score | Area under ROC | MCC |
|---|---|---|---|
| **NF** | 0.63 | 0.64 | 0.26 |
| **NF** + WCC + CPL + CFC | 0.68 | 0.64 | 0.14 |
| Held out feature | | | |
| CLC | 0.36 | 0.51 | 0.04 |
| EBC | 0.36 | 0.51 | 0.04 |
| DC2 | 0.36 | 0.51 | 0.04 |
| CS | 0.36 | 0.51 | 0.04 |
| SGC | 0.36 | 0.51 | 0.04 |
| DEG | 0.36 | 0.51 | 0.04 |
| DC1 | 0.36 | 0.51 | 0.04 |
| WCC | 0.36 | 0.51 | 0.04 |
| CPL | 0.36 | 0.51 | 0.05 |
| CFC | 0.36 | 0.51 | 0.04 |

Table S2: Network feature selection using NB among WCC, CPL, CFC and those in NF

| Experimental setup | F1 score | Area under ROC | MCC |
|---|---|---|---|
| **NF** | 0.70 | 0.77 | 0.41 |
| **NF** + WCC + CPL + CFC | 0.69 | 0.77 | 0.40 |
| Held out feature | | | |
| CLC | 0.50 | 0.74 | 0.48 |
| EBC | 0.50 | 0.74 | 0.48 |
| DC2 | 0.50 | 0.74 | 0.48 |
| CS | 0.47 | 0.73 | 0.45 |
| SGC | 0.31 | 0.75 | 0.27 |
| DEG | 0.46 | 0.73 | 0.42 |
| DC1 | 0.50 | 0.74 | 0.48 |
| WCC | 0.50 | 0.74 | 0.48 |
| CPL | 0.46 | 0.73 | 0.43 |
| CFC | 0.50 | 0.74 | 0.48 |

Table S3: Network feature selection using SVM among WCC, CPL, CFC and those in NF

| Experimental setup | F1 score | Area under ROC | MCC |
|---|---|---|---|
| **NF** | 0.74 | 0.82 | 0.50 |
| **NF** + WCC + CPL + CFC | 0.74 | 0.81 | 0.47 |
| Held out feature | | | |
| CLC | 0.54 | 0.74 | 0.50 |
| EBC | 0.54 | 0.74 | 0.50 |
| DC2 | 0.51 | 0.74 | 0.48 |
| CS | 0.53 | 0.74 | 0.50 |
| SGC | 0.53 | 0.74 | 0.50 |
| DEG | 0.50 | 0.73 | 0.47 |
| DC1 | 0.51 | 0.74 | 0.47 |
| WCC | 0.54 | 0.74 | 0.52 |
| CPL | 0.52 | 0.75 | 0.49 |
| CFC | 0.54 | 0.75 | 0.51 |

Table S4: Network feature selection using ANN among WCC, CPL, CFC and those in NF

#### S2 Hold-out evaluation among features in NF

| Experimental setup | F1 score | Area under ROC | MCC |
|---|---|---|---|
| NF | 0.63 | 0.64 | 0.26 |
| Held out feature | | | |
| CLC | 0.36 | 0.51 | 0.04 |
| EBC | 0.34 | 0.51 | 0.04 |
| DC2 | 0.36 | 0.52 | 0.04 |
| CS | 0.32 | 0.51 | 0.04 |
| SGC | 0.36 | 0.51 | 0.04 |
| DEG | 0.35 | 0.51 | 0.04 |
| DC1 | 0.36 | 0.51 | 0.04 |

Table S5: Importance of features in NF using hold out experiments for NB classifier

| Experimental setup | F1 score | Area under ROC | MCC |
|---|---|---|---|
| NF | 0.70 | 0.77 | 0.41 |
| Held out feature | | | |
| CLC | 0.50 | 0.74 | 0.48 |
| EBC | 0.50 | 0.74 | 0.48 |
| DC2 | 0.50 | 0.74 | 0.48 |
| CS | 0.47 | 0.73 | 0.45 |
| SGC | 0.41 | 0.75 | 0.37 |
| DEG | 0.46 | 0.73 | 0.42 |
| DC1 | 0.58 | 0.75 | 0.49 |

Table S6: Importance of features in NF using hold out experiments for SVM classifier

| Experimental setup | F1 score | Area under ROC | MCC |
|---|---|---|---|
| NF | 0.74 | 0.82 | 0.50 |
| Held out feature | | | |
| CLC | 0.54 | 0.74 | 0.46 |
| EBC | 0.54 | 0.74 | 0.46 |
| DC2 | 0.55 | 0.74 | 0.47 |
| CS | 0.55 | 0.74 | 0.46 |
| SGC | 0.55 | 0.74 | 0.48 |
| DEG | 0.60 | 0.76 | 0.49 |
| DC1 | 0.60 | 0.76 | 0.48 |

Table S7: Importance of features in NF using hold out experiments for ANN classifier

### S3 Evaluation of sequence features

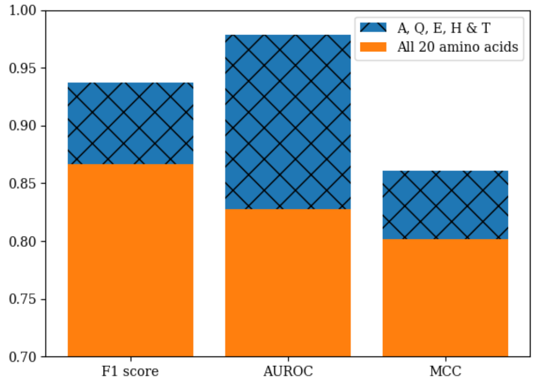


Figure S1: Comparison of RF classification using 5 chosen amino acids *vs* all 20 amino acids as features.

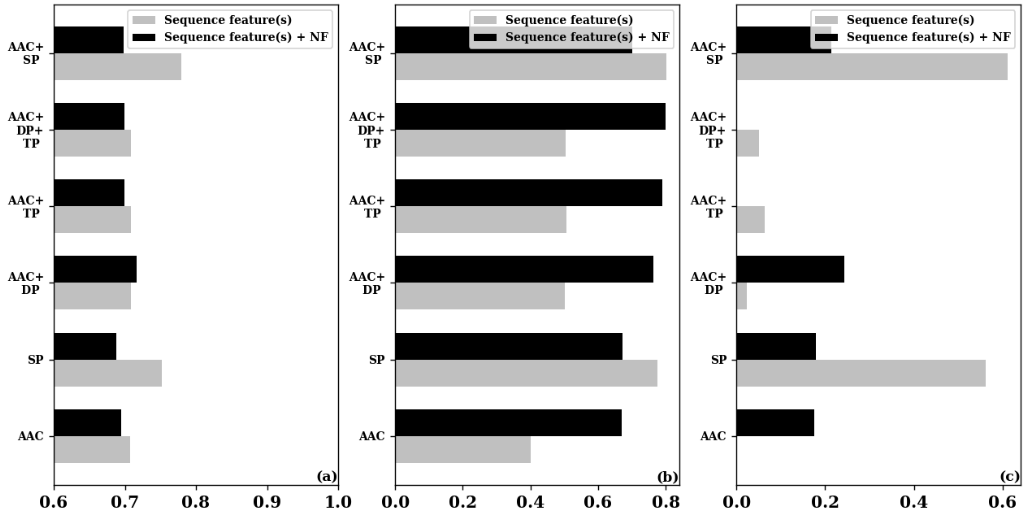


Figure S2: Comparison of classification using different combinations of network and sequence features using NB: (a) F1 score (b) Area under ROC curve (c) MCC.

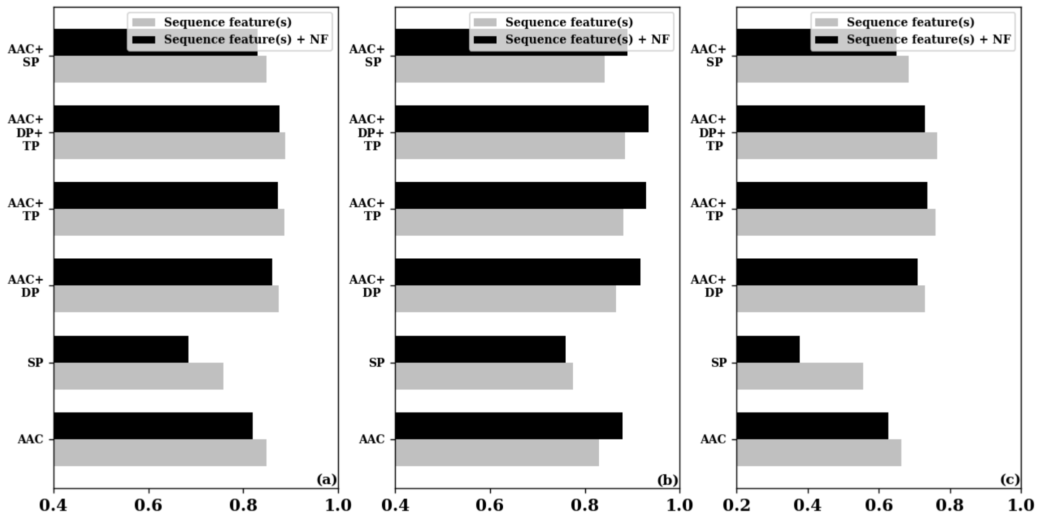


Figure S3: Comparison of classification using proposed network and sequence features using SVM: (a) F1 score (b) Area under ROC curve (c) MCC.

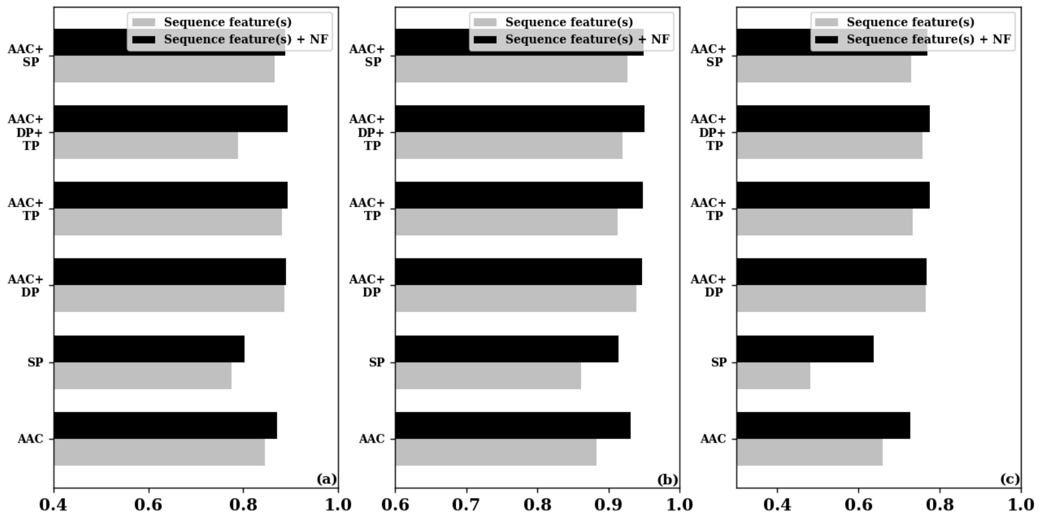


Figure S4: Comparison of classification using proposed network and sequence features using ANN: (a) F1 score (b) Area under ROC curve (c) MCC.

| Amino acid sequences | Importance(%) |
|---|---|
| gln | 13.37 |
| ile-ile | 11.13 |
| ala | 5.82 |
| met-asp | 5.77 |
| asp-glu-ile | 5.58 |
| leu-val-leu | 4.81 |
| thr | 4.57 |
| tyr-ile | 4.20 |
| glu | 3.74 |
| met-asn | 3.19 |
| his | 3.02 |
| phe-arg | 2.72 |
| gly-met | 2.68 |
| leu-arg-glu | 2.64 |
| ala-glu-glu | 2.59 |
| asn-tyr | 2.53 |
| asp-ile-ile | 2.47 |
| asn-arg | 2.43 |
| asn-val | 2.38 |
| asp-ala-leu | 2.25 |
| glu-phe | 2.11 |
| phe-ile | 1.75 |
| his-gly | 1.62 |
| asn-glu | 1.53 |
| asp-asn | 1.39 |
| met-phe | 1.27 |
| ala-ala-ala | 1.02 |
| arg-met | 0.81 |
| his-pro | 0.63 |

Table S8: Feature importance for 29 amino acid sequences using RF.
